# Proteomic analysis reveals distinct germination strategies in polymorphic fruits of *Haloxylon ammodendron*

**DOI:** 10.1101/2025.06.17.660099

**Authors:** Ziyi Wang, Weizhi Chen, Xianhua Zhang, Ze Wang, Lamei Jiang, Amanula Yimingniyazi, Cai Ren

## Abstract

Fruit polymorphism, the production of multiple fruit morphotypes within a species, is an adaptive bet-hedging strategy in variable environments. However, researches about perennial plant and the adaptive mechanisms are not well understood. *Haloxylon ammodendron*, a constructive desert shrub, exhibits three fruit morphotypes: YY (yellow wings with yellow pericarp), YP (yellow wings with pink pericarp), and PP (pink wings with pink pericarp). We investigated their ecophysiological and molecular mechanisms through germination assays under salt and drought stress, combined with proteomic analysis. YP consistently showed the highest germination percentage (GP) and germination rate index (GRI) under stress, while PP displayed well germination success under low salinity and well-watered conditions (GP=32.7%, 36.7%; GRI=0.018, 0.020), but significantly impaired viability under stress (GP=12.7%, 12.0%; GRI=0.006, 0.006). Proteomics identified 721 differentially expressed proteins (DEPs), with the most (662) between YP and PP, linked to stress response and germination. YP’s high abundance of stress-resistant proteins enabled rapid germination, whereas PP’s delayed germination aligns with a persistent seed bank strategy. This polymorphism promotes niche differentiation: YP ensures quick colonization, PP enhances long-term resilience, and YY offers an intermediate strategy. Our findings reveal molecular-ecological adaptations in *H. ammodendron*, aiding targeted germplasm use for desert restoration.

## 1. Introduction

In salt marsh and desertified extreme heterogeneous environments, elevated soil salinity inhibits seed germination in most plant species. Germination only occurs when natural precipitation temporarily alleviates this stress by reducing soil salt concentration (Huang et al., 2003; Lu and Zhang, 2015). Rainfall not only modulates salinity but also serves as one of the most critical environmental factors governing seed germination (Huang, 2016). When soil moisture fails below germination thresholds, seed dormancy persists a phenomenon termed drought induced dormancy (Yan, 2001). Desert exhibit highly sporadic and unpredictable precipitation patterns. To capitalize on ephemeral water availability, many desert plants have evolved rapid germination strategies (Han et al., 2011; Liu et al., 2019). However, the early stages of germination and seedling establishment are particularly susceptible to soil water deficit. Accelerated germination provides an adaptive advantage only when subsequent moisture supports seedling development until drought tolerance is achieved. Conversely, under conditions of insufficient precipitation, delayed or asynchronous germination becomes more advantageous (Grigg et al., 2008; Hartman et al., 2012). Through prolonged evolutionary adaptation, plants have developed diverse germination traits and survival strategies to respond to fluctuating environmental stresses, thereby optimizing the spatiotemporal regulation of germination and seedling establishment (Ungar and Khan, 2001). Fruit polymorphism represents a key adaptive strategy in extreme environments such as salt marsh and desert environments. By producing morphologically and ecologically heterogeneous diaspores, plants increase offspring survival probability in highly variable habitats, enhancing colonization success in unpredictable environments (Orlovsky et al., 2016; Hasanuzzaman et al., 2019).

Fruit polymorphism describes the phenomenon whereby a single species produces morphologically distinct fruit types within a shared habitat (genetic polymorphism), or exhibits differential fruit morphologies and behavioral traits across distinct positions on individual plants (somatic polymorphism) (Harper, 1970; Imbert, 2002). This adaptive strategy confers three primary ecological advantages: alleviation of density-dependent fitness costs in offspring (Venable et al., 1995), minimization of intraspecific competition among progeny (Mandák, 1997), and adoption of a bet hedging strategy to cope with spatiotemporal environmental heterogeneity (Venable, 1985). These adaptive features make it particularly valuable for investigating plant ecological adaptation strategies (Baskin et al., 2014; Sendek et al., 2015). Contemporary researches on fruit polymorphism have achieved significant progress in understanding ephemeral plant systems, particularly in characterizing polymorphic traits and their associated ecological behaviors including seed dispersal and germination strategies (Venable, 2007; Wang et al., 2010; Baskin and Baskin, 2023). However, significant knowledge gaps persist in two key areas: perennial plant adaptive strategies and the molecular mechanisms governing ecological behavioral divergence (Bhatt et al., 2017; Bhatt et al., 2019; El-Keblawy et al., 2013). Proteomic methodologies provide powerful tools to address these limitations through comparative protein expression analysis across different fruits types. Functioning as both structural components and functional regulators, proteins participate in nearly all essential physiological processes. Moreover, as one of three major seed storage reserves (along with carbohydrates and lipids), proteins are pivotal for seed vigor and germination (Hu, 2006). The proteome, defined as the complete set of proteins expressed at cellular, tissue, and organismal levels, serves as a key analytical framework for investigating seed vigor, germination, and seedling establishment mechanisms (Wilkins et al., 1996; Wang et al., 2023). The seed appears to be well prepared to mobilize the major classes of reserves (proteins, triglycerides, phytate, and starch) during germination, indicating that the preparation of the seed for germination is mainly achieved during its maturation on the mother plant. Furthermore, proteome analyses have revealed several pathways that can contribute to seed vigor, an important agronomic trait defined as the potential to produce vigorous seedlings, such as glycine betaine accumulation in seeds (Catusse et al., 2008; Nguyen et al., 2015; Sajeev et al., 2024). Consequently, germination performance can be mechanistically explained by the seed’s protein profile.

*Haloxylon ammodendron* (Amaranthaceae), a shrub, serves as a constructive species in desert vegetation communities. In China, it is primarily distributed in the Gurbantunggut Desert of the Junggar Basin, where it forms constructive communities providing critical habitat for nearly 200 desert plant species (Xiao et al., 2019). These communities make these ecosystems vital for biodiversity conservation in arid regions. Thus, *H. ammodendron* plays an irreplaceable ecological role in windbreak, desertification mitigation, and maintenance of regional ecological balance (Jia et al., 2004). This shrub survives under extreme environmental conditions characterized by: extreme summer air temperatures exceeding 40□, annual evaporation rates surpassing 2000 mm, mean annual precipitation below 150 mm, soil salinity up to 2% in highly saline areas (Liu et al., 2009). To cope with these challenging conditions, *H. ammodendron* has developed a fruit polymorphism strategy as an ecological adaptation. The species produces plants with three distinct fruit morphotypes within the same habitat characterized by different wing and pericarp color combinations (genetic polymorphism): yellow wings with yellow pericarp (YY), yellow wings with pink pericarp (YP), and pink wings with pink pericarp (PP) (Fig. 1). Notably, these polymorphic fruits exhibit differential germination responses to salinity and moisture gradients, representing a bet-hedging strategy to maximize reproductive success across spatially and temporally heterogeneous conditions (Wang et al., 2025).

**Fig. 1.**
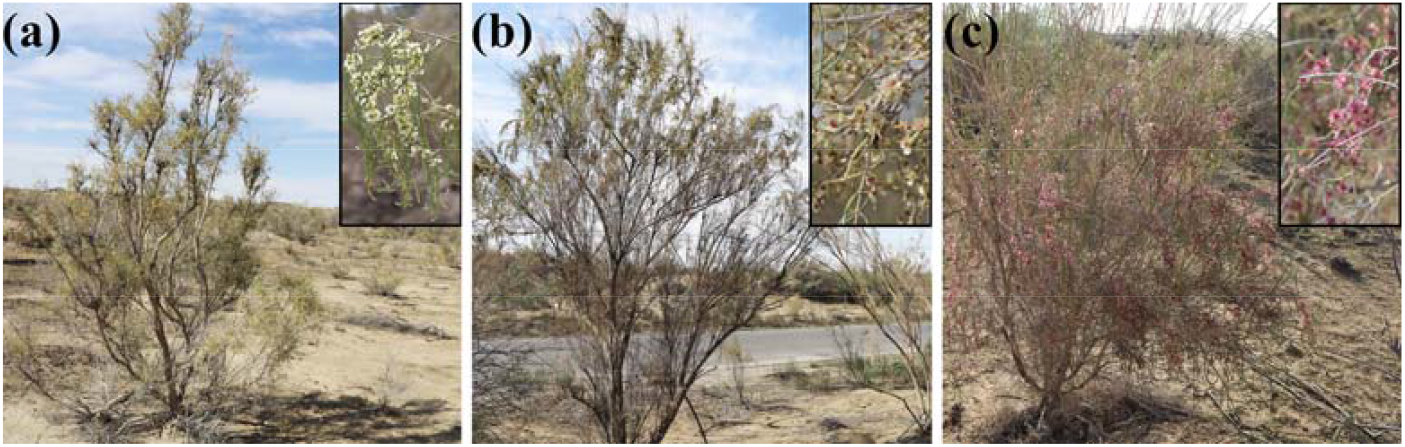
Habitat and fruit polymorphism in *Haloxylon ammodendron*. (a) YY type plant: fruit with yellow wings and yellow pericarp; (b) YP type plant: fruit with yellow wings and pink pericarp; (c) PP type plant: fruit with pink wings and pink pericarp.

Salt and drought are the main environmental factors affecting plant reproduction, dormancy, and germination. Although the three fruit types of *H. ammodendron* exhibit differential responses to these stresses, the underlying protein-level mechanisms remain unclear. This study aims to: analyze the stress response variations among the three fruit types and their proteomic basis through controlled greenhouse germination experiments. The findings will provide a new theoretical foundation for comprehensively understanding plant adaptation mechanisms in desert environments.

## 2. Materials and method

### 2.1. Study area and sampling

The study area is located in the desert region along the southern margin of the Junggar Basin (44.37°N, 87.96°E), one of the primary distribution zones for *H. ammodendron* populations in Northwest of China. Characterized by a typical continental arid to semi-arid climate, the Junggar Basin receives an average annual precipitation of less than 150 mm. Soil salinity in the most saline areas exceeds 10 g kg^-1^, with hardened white salt crusts observed in localized regions (Ma et al., 2018). Additionally, intersecting sand dunes generate pronounced spatiotemporal heterogeneity in soil salinity and moisture conditions, creating a highly unpredictable environment that limits seedling survival and population regeneration of *H. ammodendron* (Wang et al., 2025). During the fruit maturation period in November 2023, three types of intact fruits were collected, cleaned of impurities, and stored at -20°C for subsequent experiments.

### 2.2. Germination responses of polymorphic fruits to varying salinity and water availability

This study employed a single-factor experimental design to compare the germination characteristics of three fruit types under varying salinity and moisture conditions. Thirty seeds of each fruit type were individually sown in pots (30 cm in diameter and depth) filled with native soil, with six replicates per treatment. Four salinity treatments were established with soil NaCl concentrations of 0, 7, 13, and 16 g kg^-1^ (Agricultural Bureau and Soil Survey of Office of Uygur Autonomous region of Xinjiang, 1996). The pots were irrigated daily to maintain field capacity (20% soil gravimetric water content). Additionally, three moisture regimes were applied by watering to field capacity at intervals of 1, 3, and 6 days. Seedling emergence (defined as cotyledon appearance) was recorded daily for 28-day. Germination percentage and germination rate index were calculated using established formulas (Hsu et al., 1985):

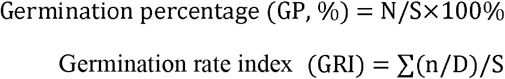

Where, N = total number of germinated seeds, S = total number of seeds tested, n = number of seeds germinated on day D, D = number of days since sowing (the maximum possible GRI value is 100).

### 2.3. Proteomic sequencing and bioinformatics analysis

The total proteins from three types of fruits were extracted using the trichloroacetic acid (TCA)/acetone precipitation method, followed by sample preparation. Protein concentration was determined using the Bradford protein assay kit. Subsequently, the proteins underwent reduction, alkylation, enzymatic digestion, and labeling. The resulting peptides were separated by high-performance liquid chromatography (HPLC) and analyzed by mass spectrometry using an Orbitrap Fusion™ Tribrid™ system (Thermo Fisher Scientific). For proteomic data analysis, the database was derived from transcriptome sequencing data (Uniprot protein database). The screening criteria for total proteins were set at a false discovery rate (FDR) of < 1%. Differentially expressed proteins were identified based on the following criteria: a fold change ≥2.0, the presence of at least one unique peptide, and a statistically significant difference (P < 0.05) as determined by the t-test algorithm.

### 2.4. Date analysis

The raw data were processed and visualized using Excel 2016 (Microsoft, Redmond, Washington, USA, 2016), and Origin 2021 (OriginLab, Northampton, Massachusetts, USA, 2021). Statistical analysis was performed using SPSS 25 (IBM SPSS, Chicago, Illinois, USA, 2017). Prior to analysis, the Shapiro-Wilk test was used to assess normality, and Levene’s test (P > 0.05) was applied to verify homogeneity of variance. If the data met the assumptions of normality and homoscedasticity, one-way ANOVA followed by the LSD (Least Significant Difference) post hoc test (P < 0.05) was conducted. If the assumption of homogeneity of variance was violated, Tamhane’s T2 test (a non-parametric alternative) was employed instead.

## 3. Results

### 3.1. Differential germination responses of polymorphic fruits to varying salinity and water availability

Germination characteristics of polymorphic fruits exhibited significant variation in response to differential salinity and moisture regimes (Fig. 2). Notably, YP displayed consistently superior germination performance, with significantly elevated GP and GRI values across all experimental salinity gradients compared to other fruit types. At reduced NaCl concentrations (0-7 g kg^-1^), PP manifested significantly enhanced germination parameters relative to YY. Conversely, under elevated NaCl stress (13-16 g kg^-1^), a marked inversion of this pattern occurred, with PP demonstrating significantly depressed GP and GI measurements compared to YY (Fig. 2a,b).

**Fig. 2.**
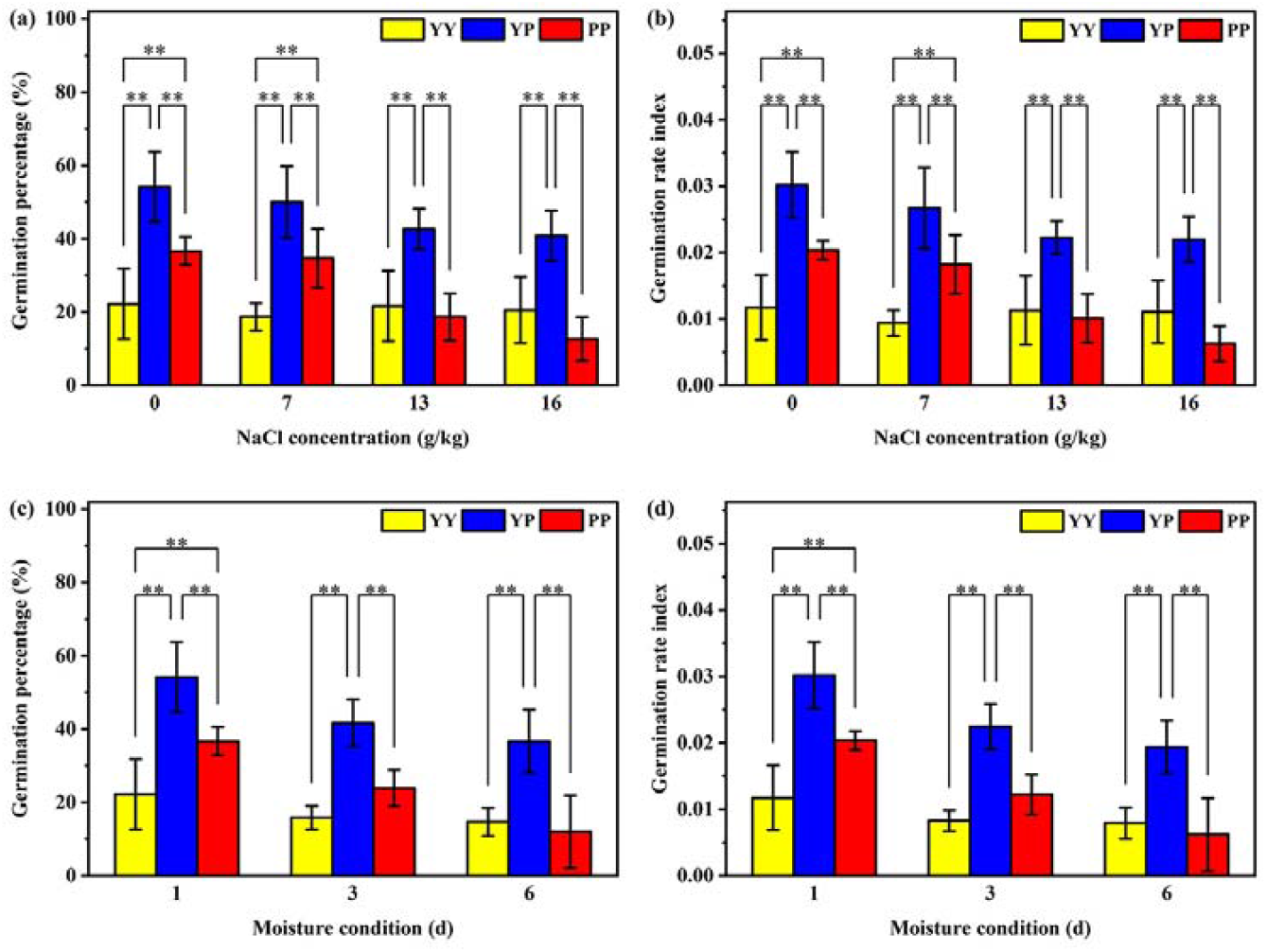
Comparative analysis of germination characteristics among three fruit morphotypes of *Haloxylon ammodendron* under differential salinity and moisture conditions. (a) Germination percentage under varying salinity treatments; (b) Germination rate index under varying salinity treatments; (c) Germination percentage under varying watering regimes; (d) Germination rate index under varying watering regimes. Data represent mean values ± standard deviation (SD). Double asterisks (**) indicate statistically significant differences (P < 0.01) between two morphotypes within the same treatment.

Across all moisture regimes, YP consistently maintained superior germination performance, exhibiting significantly higher GP and GI values compared to others’ (Fig. 2c,d). Under daily watering conditions, PP demonstrated significantly enhanced germination parameters relative to YY. Conversely, when subjected to 7-day watering intervals, PP showed markedly reduced GP and GI values compared to YY. The combined results indicate that YP possesses strong tolerance to both salinity and drought stress, while PP shows marked sensitivity to these conditions.

### 3.2. Differential protein identification and GO_BP pathway enrichment analysis

A total of 3,080 proteins were identified across the three fruit morphotypes. Principal component analysis (PCA) demonstrated strong biological reproducibility among replicates while revealing significant proteomic divergence among them (Fig. 3a). Hierarchical clustering analysis further indicated the most pronounced differential protein expression between YP and PP (Fig. 3b).

**Fig. 3.**
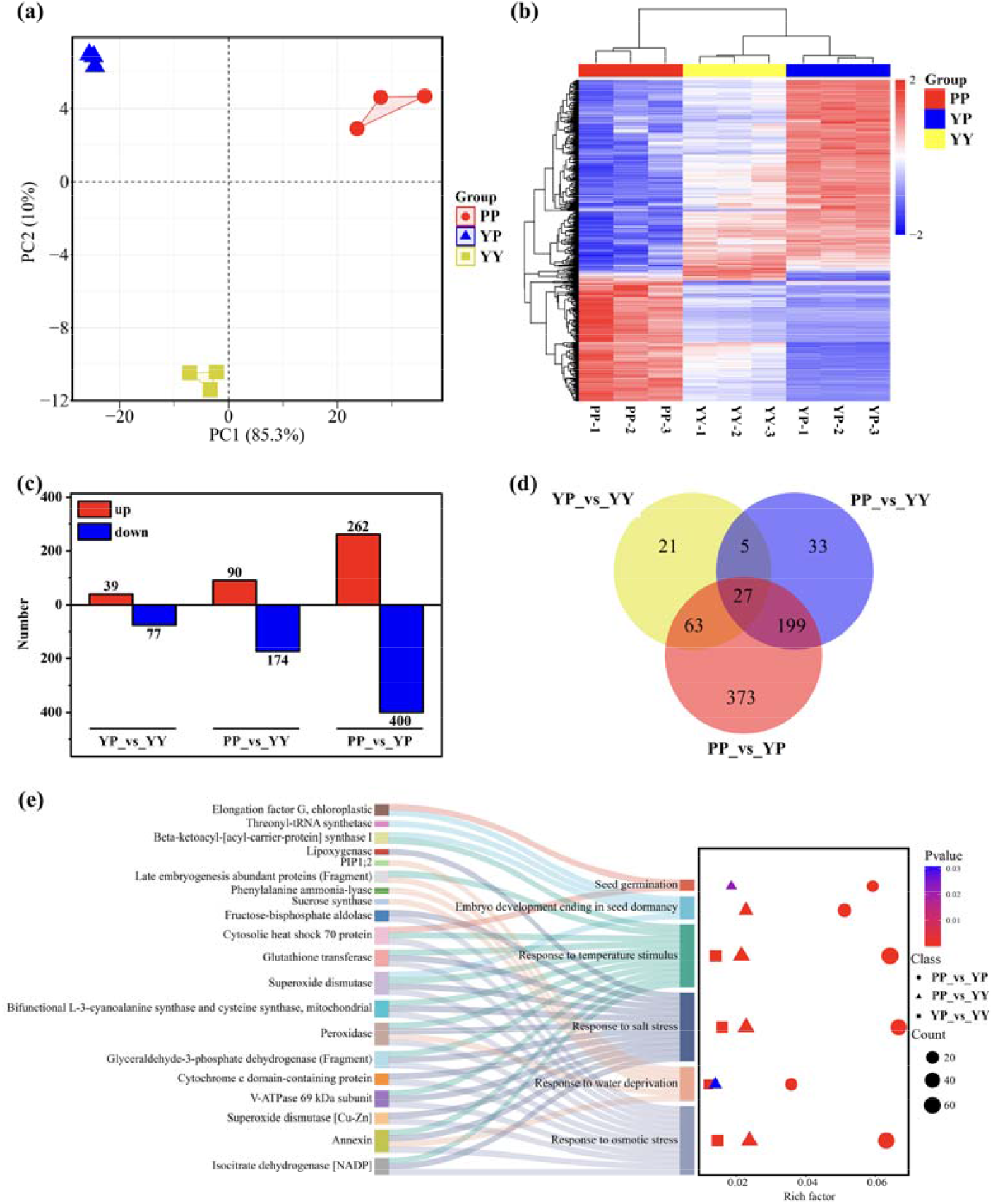
Differential protein identification and GO_BP pathway enrichment analysis in polymorphic fruits of *Haloxylon ammodendron*. (a) Principal component analysis (PCA) of polymorphic fruit. (b) Hierarchical clustering of polymorphic fruit proteomes. (c) Venn diagram of differentially expressed proteins (DEPs) among polymorphic fruit. (d) Shared and unique DEPs between polymorphic fruit. (e) Significantly enriched GO biological processes (FDR<0.05) for DEPs.

Using stringent identification criteria (FDR < 1%, fold-change ≥ 2, ≥1 unique peptide, and t-test P < 0.05), we identified 721 differentially expressed proteins (DEPs). Comparative analyses revealed: 116 DEPs between YP and YY, 264 DEPs between PP and YY, 662 DEPs between PP and YP (Fig. 3c). Notably, PP vs YP exhibited the greatest proteomic divergence, with 373 DEPs being uniquely specific to this comparison (Fig. 3d).

GO_BP enrichment analysis demonstrated significant association of DEPs in *Haloxylon* polymorphic fruits with six key biological processes (Fig. 3e), predominantly including seed dormancy regulation, germination control, and environmental stress response.

### 3.3. Proteomic analysis of differential proteins expression associated with germination traits under abiotic atress

Comparative proteomic analysis revealed 98 DAPs associated with germination-related and stress-responsive pathways across the three fruit morphotypes (Supplementary Table. 1). Notably, 20 of these DAPs represent previously documented regulators of seed germination and environmental adaptation (Fig. 4a).

**Fig. 4.**
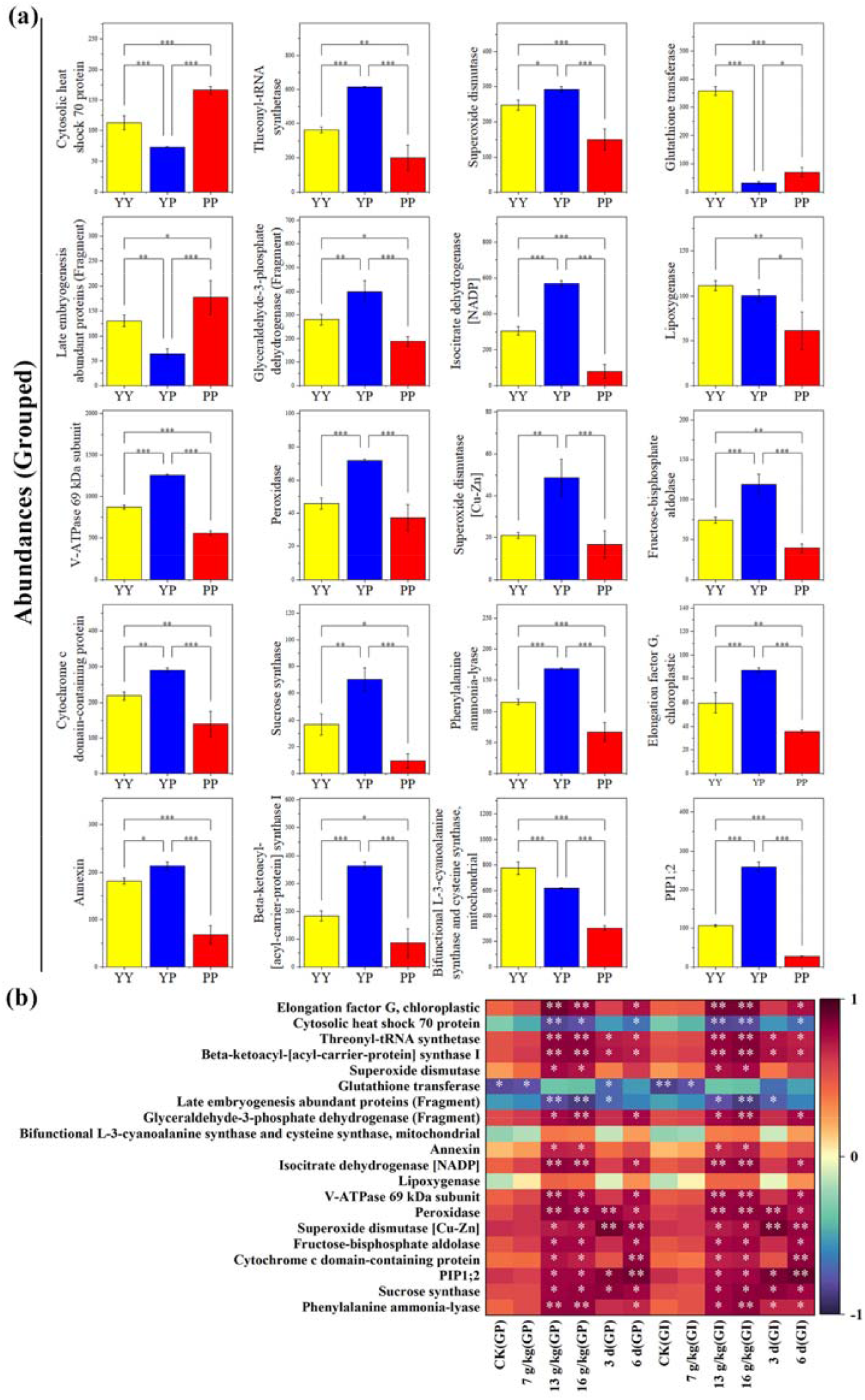
Analyses of differential protein abundance and its correlation with germination traits under stress conditions. (a) Comparative analysis of protein abundance profiles across three fruit morphotypes. Values represent normalized relative abundance (mean ± SD). Asterisks denote statistically significant intergroup differences (* P < 0.05) determined by ANOVA with post-hoc Tukey test. (b) Pearson correlation matrix between differentially accumulated proteins (DAPs) and germination parameters (germination percentage and germination index). Statistical significance of correlation coefficients is indicated by asterisks (* P < 0.05, ** P < 0.01, *** P < 0.001), with Benjamini-Hochberg correction for multiple comparisons.

Statistical evaluation of protein abundance patterns against germination metrics demonstrated significant positive correlations (n = 15) and negative correlations (n=3) between specific DAPs and germination parameters (percentage and index) in polymorphic fruits under differential stress regimes (Fig. 4b).

## 4. Discussion

### 4.1. Differential germination strategies in polymorphic fruits

Plant species exhibiting fruit polymorphism predominantly inhabit extreme environments characterized by deserts, saline marshes, and gravel substrates with elevated soil salinity and limited water availability. Both salinity and water deficit serve as critical environmental factors regulating seed dormancy and germination kinetics. Our experimental results demonstrated that under severe saline stress (16 g kg^-1^ NaCl) and prolonged drought (6 days), *H. ammodendron*’s YP fruit exhibited superior germination performance (in both GP and GRI), while the PP displayed significantly reduced germination capacity. These findings align with previous investigations on seed polymorphism, where germination differences among polymorphic diaspores under controlled conditions are well documented. For instance, *Atriplex sagittata* produces dimorphic seeds, with larger brown seeds demonstrating higher germination percentage than smaller black seeds (Mandák and Pyek, 2001). Similarly, in *Salsola splendens*, both black and brown polymorphic seeds generate viable progeny under 0.4 mol L^-1^ NaCl (approximating natural habitat salinity), though brown-seeded progeny exhibit reduced biomass accumulation at sub-optimal salinity (0.2 mol L^-1^ NaCl) (Redondo-Gómez et al., 2008). Ecologically significant is *Zygophyllum xanthoxylum*’s germination response: progressive inhibition occurs below -0.3 MPa water potential, with complete germination cessation at -1.5 MPa. These thresholds reveal critical thresholds for seedling establishment in arid environments (Zeng et al., 2004).

Experimental investigations revealed distinct adaptive strategies between fruit morphotypes under salt and drought stress conditions. YP type fruits displayed the highest germination capacity and seedling emergence rates under salt and drought stress conditions. However, this apparent competitive advantage was offset by significantly reduced seedling survivorship, creating substantial resource allocation inefficiencies during early establishment phases. Conversely, PP type fruits employed a conservative strategy characterized by stress-responsive germination suppression, thereby maintaining fruit viability and establishing a transient soil seed bank. Following precipitation events (either snowmelt or rainfall) that reduced surface soil salinity and enhanced moisture availability, these dormant seeds initiated germination and emergence, leading to significantly enhanced seedling survival rates (Zhumabekova et al., 2020).

### 4.2. Comparative proteomic analysis of polymorphic fruits

Comparative proteomic profiling and cluster analysis of three fruit morphotypes revealed significant differential protein accumulation between PP type and YP type fruits, with 662 proteins exhibiting distinct abundance patterns. Hierarchical clustering further elucidated both similarities and divergences in protein expression profiles among the three fruit types. Notably, PP type and YP type fruits displayed the most pronounced proteomic divergence, while YY type fruits exhibited intermediate characteristics. These proteome-level variations likely constitute a fundamental molecular mechanism underlying their distinct dormancy patterns, germination behaviors, and stress response strategies.

Functional enrichment analysis of differentially expressed proteins among the three fruit morphotypes revealed significant enrichment in several key biological pathways. Notably, these proteins were predominantly enriched in pathways related to seed germination, drought response, and salt stress adaptation. The presence of these differential proteins likely contributes to the distinct germination regulation mechanisms and environmental adaptation strategies observed among the morphotypes. Particularly under the severe saline and arid conditions characteristic of desert environments, these differentially expressed proteins may mediate divergent stress response strategies through distinct regulatory mechanisms (Chen et al., 2022).

### 4.3. Proteomic insights into stress-responsive germination mechanisms of polymorphic fruits

Proteomic comparison of polymorphic fruit identified 98 co-accumulated proteins functionally linked to both germination processes and environmental stress responses. Notably, 20 of these proteins represent well-characterized regulators previously implicated in germination control and stress adaptation mechanisms. Statistical analysis of germination parameters under stress conditions revealed distinct correlation patterns: 15 proteins showed significant positive correlations with both germination percentage and germination index, 3 proteins displayed significant negative correlations, while two proteins showed no statistically significant association.

Proteomic analysis identified several stress-responsive proteins exhibiting both positive correlations with germination parameters under abiotic stress conditions, and significantly higher abundance in YP type versus PP type fruits. These molecular determinants of germination efficiency included: Chloroplastic elongation factor G (*CPEFG*), *Arabidopsis* mutants exhibit delayed germination, indicating its essential role in translation initiation during early germination events (Albrecht et al., 2006); Threonyl-tRNA synthetase (*EMB2761*), mediates threonine-tRNA charging, thereby maintaining the translational capacity required for germination-associated protein synthesis (Barai and Chen, 2024); β-Ketoacyl-ACP synthase I (*KAS2*), catalyzes the initial elongation step in de novo fatty acid biosynthesis (C4→C16), providing lipid reserves for energy mobilization during germination (Yang et al., 2016); Isocitrate dehydrogenase [NADP] (*CICDH*), biochemical studies in *Gossypium hirsutum* demonstrate a direct correlation between ICDH activity and germination kinetics (Tsaniklidis et al., 2015); Cytochrome c domain protein (*CYC11*), *Arabidopsis* cyc11 knockouts show delayed radicle emergence, implicating mitochondrial electron transport in germination vigor (Racca et al., 2022); Annexin D5 (*ANN5*), structural homolog of NnANN1 from *Nelumbo nucifera*, which play crucial roles in seed vigor (Chu et al., 2012); Class III peroxidase (*PRX3*), modulates ABA/GA balance through phytohormone metabolism, positively regulates germination potential in cowpea (*Vigna unguiculata*) and wheat (*Triticum aestivum*) (Gao et al., 2024).

Proteomic analysis identified several stress-responsive proteins with significantly higher abundance in YP type fruits compared to PP-type, providing a molecular basis for their enhanced germination speed and stress resistance: Superoxide dismutase [Cu-Zn] (*MSD1/CSD2*), demonstrates ROS-scavenging capacity in *Nitraria roborowskii*, and enhances salt tolerance. Over expression of *Potentilla CSD2* in *Arabidopsis* improves seed germination under saline stress (Gill et al., 2012); Vacuolar-type H^+^-ATPase subunit A (*V-ATPase*), over expression of the wheat *TaVB* gene (encoding V-ATPase subunit B) in *Arabidopsis* enhances salt tolerance by maintaining ion homeostasis (Wang et al., 2011); Glyceraldehyde-3-phosphate dehydrogenase (Fragment) (*GAPC2*), found to be drought-responsive in the moss *Racomitrium japonicum*, suggesting a role in stress adaptation (Zhang et al., 2014); Plasma membrane intrinsic protein PIP1;2 (*PIP1;2*), indicates its involvement in rapid water uptake during drought and salt stress, promoting germination in *Brassica napus* and *Spinacia oleracea* (Chen et al., 2013); Phenylalanine ammonia-lyase (*PAL1*), over expression of *GmPAL1.1* in *Arabidopsis* enhances tolerance to both salt and drought stress, likely through phenylpropanoid pathway activation, negative regulator of germination under stress (Wen et al., 2008); Late embryogenesis abundant protein (Fragment) (*SAG21*), shows negative correlation with germination rate/speed under stress (Zhang et al., 2013); LEA proteins accumulate under osmotic stress, and their higher abundance in PP type fruits may indicate greater stress susceptibility and suppressed germination capacity.

Proteomic analysis identified significant enrichment of temperature-stress related proteins, revealing distinct adaptation strategies among fruit morphotypes: Fructose-bisphosphate aldolase (*FBA6*), shows higher abundance in YP type fruits, enhances seed tolerance to both low and high temperature stress during germination in *Solanum lycopersicum* (Cai et al., 2016); Sucrose synthase (*SUS3*), overexpressing tomato lines exhibit lower malondialdehyde (MDA) accumulation under cold stress, higher proline and soluble sugar content, maintain superoxide dismutase (SOD)/catalase (CAT) activity, associated with improved cold-stress germination capacity (Li et al., 2024); Cytosolic heat shock 70 protein (*HSP70/MED37E*), predominantly accumulates in PP type fruits, Its *Arabidopsis* ortholog modulates cold response through ABA biosynthesis induction, which inhibits seed germination (Ashraf et al., 2021). Consistent with these proteomic patterns, YP type fruits of *H. ammodendron* demonstrated accelerated germination under low temperatures (Wang et al., 2025). The differential protein profiles directly correlated with observed thermo-tolerance phenotypes.

Glutathione transferase (*GSTF8*), Bifunctional L-3-cyanoalanine synthase/cysteine synthase (*CYSC1*), and lipoxygenase (*LOX2*) were predominantly expressed in YY type fruits. The first two proteins play a crucial role in enhancing seed tolerance to salt stress (Horváth et al., 2019; Yu et al., 2021), whereas LOX2 is involved in mediating seed germination under drought conditions, with its activity being markedly upregulated in response to drought stress (Chen, et al., 2018). These proteins may collectively contribute to the enhanced stress resistance capacity observed in YY type fruits.

Integrated analysis of polymorphic fruit germination traits and proteomic profiles revealed that differentially expressed proteins among polymorphic fruit types coordinately regulate ecological adaptation through multiple physiological processes, thereby establishing distinct germination and stress response strategies. In YP type fruits, the elevated abundance of stress resistance and germination promoting proteins maintained high germination percentage and rapid germination kinetics under both high salinity and severe drought stress conditions, demonstrating enhanced stress tolerance. Conversely, PP type fruits showed reduced abundance of stress responsive proteins, resulting in either complete germination inhibition or minimal germination under high salinity and drought stress. This adaptive mechanism facilitated transient soil seed bank formation, enabling avoidance of adverse environmental conditions.

## 5. Conclusion

This study reveals that *H. ammodendron* has evolved fruit polymorphism strategies as an adaptive mechanism to highly heterogeneous harsh environment. Three discrete fruit morphotypes demonstrate differential stress-response strategies. YP type fruits exhibit high abundance of stress resistance and germination promoting proteins, demonstrating strong stress tolerance with high germination rates and speed under both high salinity and severe drought stress conditions. In contrast, the PP type fruits show low abundance of stress related proteins, resulting in minimal germination, thereby forming a transient soil seed bank to avoid unfavorable conditions. The YY type fruits display intermediate characteristics (Fig. 5). The differential responses of the three fruit types to environmental stresses enable them to effectively cope with high environmental heterogeneity, thereby adopting a “bet-hedging” life history strategy to accomplish seedling establishment and population regeneration. This study elucidates the ecological adaptation mechanisms of fruit polymorphism in *H. ammodendron*, providing novel theoretical insights into plant adaptive strategies in desert environments.

**Fig. 5.**
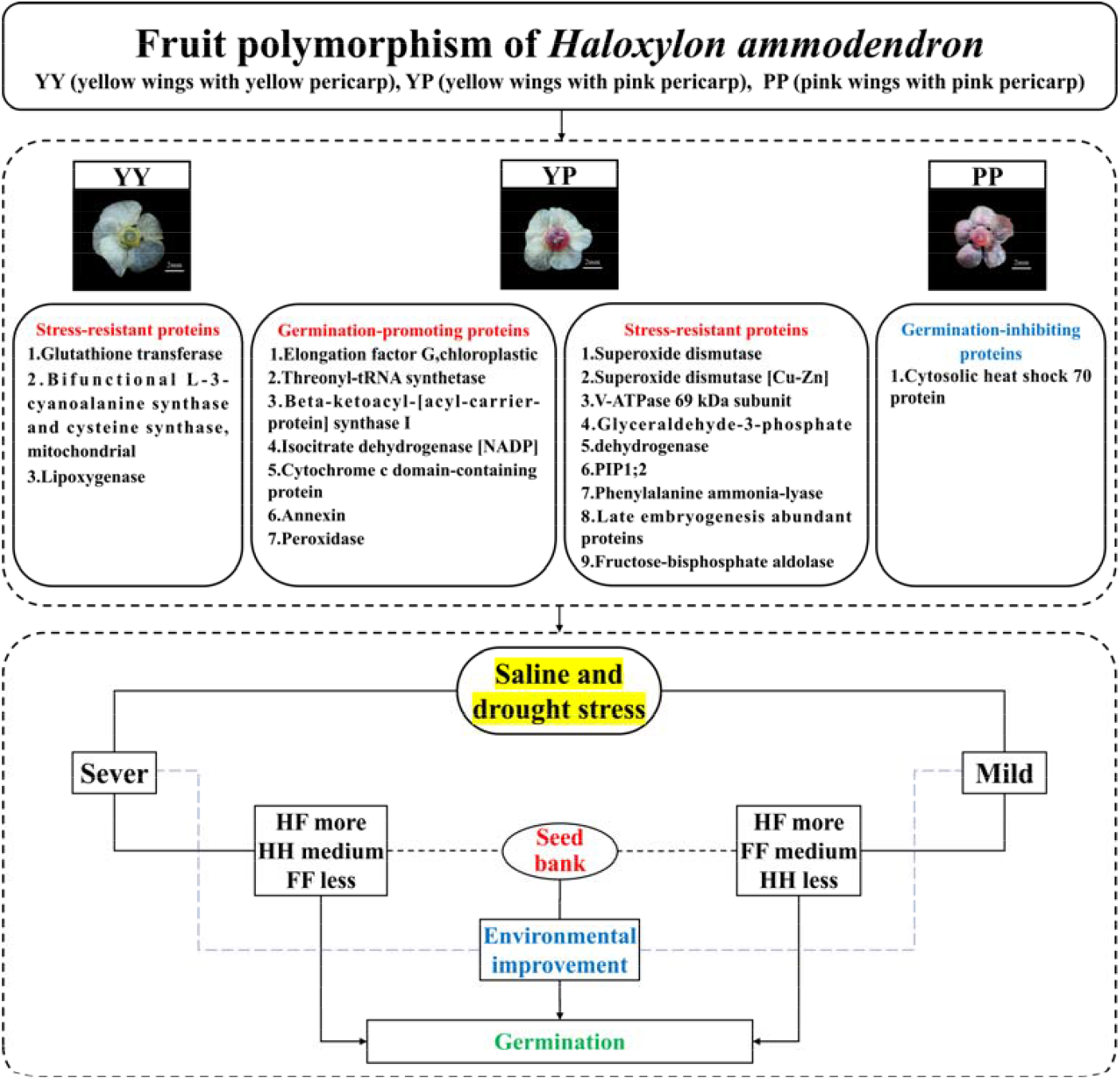
Schematic diagram of differential protein-mediated germination characteristics among three fruit morphotypes in *Haloxylon ammodendron*.

## Author contributions

Ziyi Wang, Weizhi Chen, Xianhua Zhang, Ze Wang and Cai Ren: conception and design of the experiments; Ziyi Wang, Weizhi Chen and Lamei Jiang: performance of the experiments; Amanula Yimingniyazi and Cai Ren: writing of the manuscript.

## Data availability

All data supporting the findings of this study are available within the paper and within its supplementary materials published online.

---

**Supplementary Table. 1.**
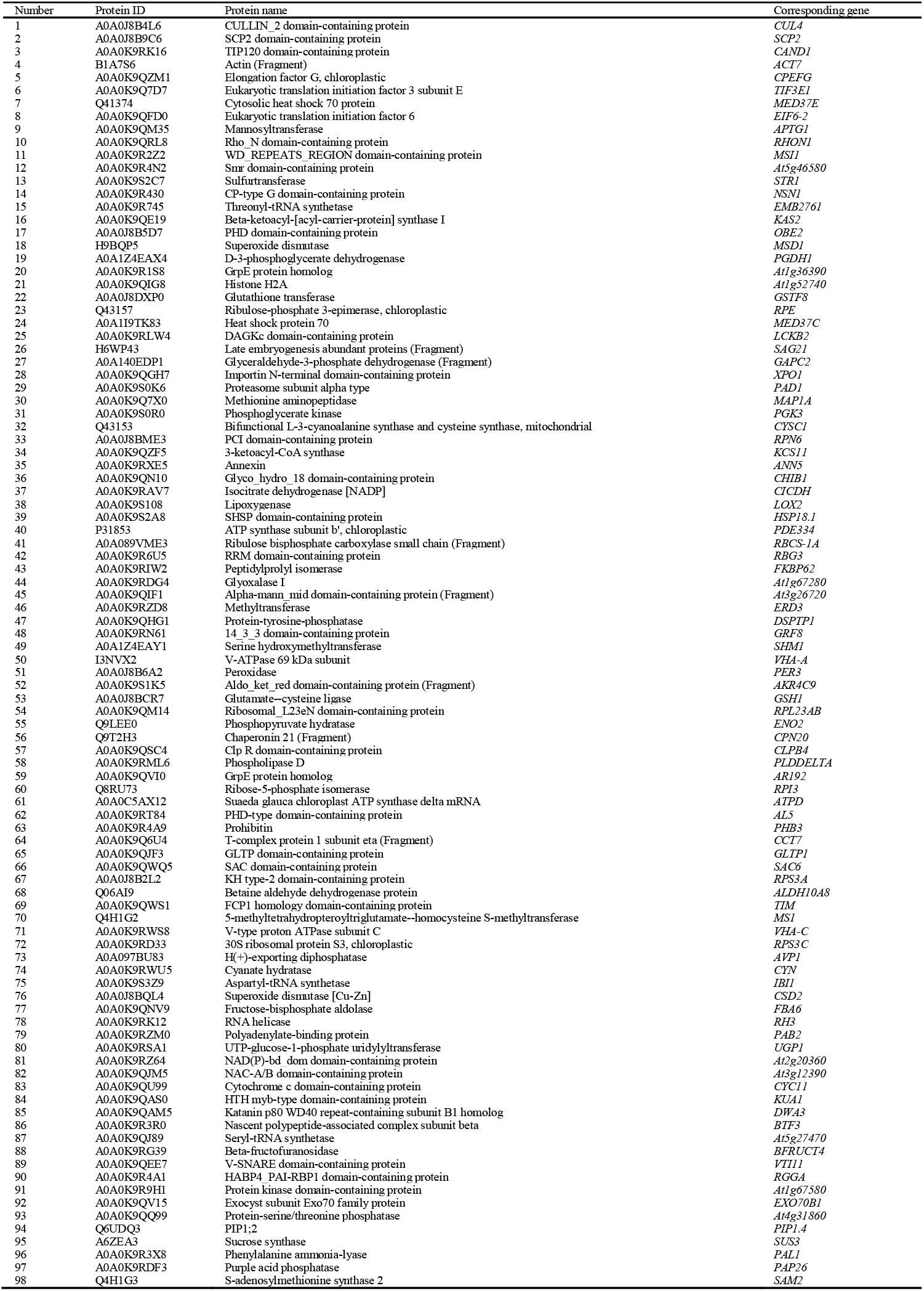
Functional proteomic profile of germination-related and stress-responsive proteins across fruit morphotypes in *Haloxylon ammodendron*.

## References

Agricultural Bureau and Soil Survey Office of Uygur Autonomous region of Xinjiang. 1996. Soil in Xinjiang. Beijing: Science Press, 51–52.

Albrecht V, Ingenfeld A, Apel K. 2006. Characterization of the snowy cotyledon 1 mutant of Arabidopsis thaliana: the impact of chloroplast elongation factor G on chloroplast development and plant vitality. Plant Molecular Biology 60, 507–518.

Ashraf M, Mao QL, Hong J, et al. 2021. HSP70-16 and VDAC3 jointly inhibit seed germination under cold stress in Arabidopsis. Plant Cell and Environment 11, 3616–3627.

Barai P, Chen J. 2024. Beyond protein synthesis: non-translational functions of threonyl-tRNA synthetases. Biochemical Society Transactions 52, 661–670.

Baskin JM, Baskin CC. 2023. A classification system for seed (diaspore) monomorphism and heteromorphism in angiosperms. Seed Science and Technology 33, 193–202.

Baskin JM, Lu JJ, Baskin CC, et al. 2014. Diaspore dispersal ability and degree of dormancy in heteromorphic species of cold deserts of northwest China: A review. Perspectives in Plant Ecology, Evolution and Systematics 16, 93–99.

Bhatt A, Bhat NR, Carón MM, et al. 2019. Dimorphic fruit colour is associated with differences in germination of Calligonum comosum. Botany 97, 263–268.

Bhatt A, Phartyal SS, Nicholas A. 2017. Ecological role of distinct fruit-wing perianth color in synchronization of seed germination in Haloxylon salicornicum. Plant Special Biology 32, 121–133.

Cai BB, Li Q, Xu YC, et al. 2016. Genome-wide analysis of the fructose 1,6-bisphosphate aldolase (FBA) gene family and functional characterization of FBA7 in tomato. Plant Physiology and Biochemistry 108, 251–265.

Catusse J, Strub JM, Job C, et al. 2008. Proteome-wide characterization of sugarbeet seed vigor and its tissue specific expression. Proceedings of the National Academy of Sciences of the United States of America 105, 10262–10267.

Chen K, Fessehaie A, Arora R. 2013. Aquaporin expression during seed osmopriming and post-priming germination in spinach. Biologia Plantarum 57, 193–198.

Chen L, Liu B, An YH, et al. 2018. Effects of phospholipase Dδ and 9-lipoxygenase on jasmonic acid synthesis and seed germination of Arabidopsis thaliana under drought stress. Chinese Journal of Ecology 37, 2627–2636.

Chen YY, Wang JC, Yao LR, et al. 2022. Combined proteomic and metabolomic analysis of the molecular mechanism underlying the response to salt stress during seed germination in barley. International Journal of Molecular Sciences 23, 10515.

Chu P, Chen HH, Zhou YL, et al. 2012. Proteomic and functional analyses of Nelumbo nucifera annexins involved in seed thermotolerance and germination vigor. Planta 235, 1271–1288.

El-Keblawy AA, Bhatt A, Gairola S. 2013. Perianth colour affects germination behaviour in wind-pollinated Salsola rubescens in Arabian deserts. Botany 92, 69–75.

Gao W, Jiang YT, Yang XH, et al. 2024. Functional analysis of a wheat class III peroxidase gene, TaPer12-3A, in seed dormancy and germination. BMC Plant Biology 24, 318.

Gill T, Dogra V, Kumar S, et al. 2012. Protein dynamics during seed germination under copper stress in Arabidopsis over-expressing Potentilla superoxide dismutase. Journal of plant research 125, 165–172.

Grigg AM, Veneklaas EJ, Lambers H. 2008. Water relations and mineral nutrition of Triodia grasses on desert dunes and interdunes. Australian Journal of Botany 56, 408–421.

Han JX, Wei Y, Yan C, et al. 2011. The vivipary characteristic of Anabasis elatior and its ecological adaptation. Acta Ecologica Sinica 31, 2662–2668.

Harper JL, Lovell PH, Moore KG. 1970. The shapes and sizes of seeds. Annual Review of Ecology Evolution and Systematics 1, 327–356.

Hartman JC, Nippert JB, Springer CJ. 2012. Ecotypic responses of switchgrass to altered precipitation. Functional Plant Biology 39, 126–136.

Hasanuzzaman M, Shabala S, Fujita M. 2019. Halophytes and climate change: adaptive mechanisms and potential uses. Boston: CABI, 104-113.

Horváth E, Bela K, Holinka B, et al. 2019. The Arabidopsis glutathione transferases, AtGSTF8 and AtGSTU19 are involved in the maintenance of root redox homeostasis affecting meristem size and salt stress sensitivity. Plant Science 283, 366–374.

Hsu FH, Nelson CJ, Matches AG. 1985. Temperature effects on germination of perennial warm-season forage grasses. Crop Science 25, 215–220.

Hu J. 2006. Seed biology. Beijing: Higher Education Press, 50.

Huang J. 2016. Accelerated dryland expansion under climate change. Nature Climate Change 6, 166–171.

Huang ZY, Zhang XS, Zheng GH, et al. 2003. Influence of light, temperature, salinity and storage on seed germination of Haloxylon ammodendron. Journal of Arid Environments 55, 453–464.

Imbert E. 2002. Ecological consequences and ontogeny of seed heteromorphism. Perspectives in Plant Ecology, Evolution and Systematics 5, 13–36.

Jia ZQ, Lu Q, Guo BG, et al. 2004. Progress in the study of psammophyte-Haloxylon. Forest Research 17, 125–132.

Li S, Wang Y, Liu Y, et al. 2024. Sucrose synthase gene SUS3 could enhance cold tolerance in tomato. Frontiers in Plant Science 14, 1324401.

Liu DJ, Liu YH, Sheng JD, et al. 2009. Salinity character of underlying horizon of Haloxylon ammodendron in the north of Xinjiang. Journal of Chinese Soil and Water Conservation 23, 47–51.

Liu GJ, Zhang XM, Lv CY, et al. 2019. Natural regeneration and maintenance ecology of Haloxylon ammodendron. Beijing: Science Press, 244–247.

Lu XT, Zhang HX. 2015. Salt tolerance during seed germination and early seedling stages of 12 halophytes. Plant and Soil 62, 229–241.

Mandák B. 1997. Seed heteromorphism and the life cycle of plants: a literature review. Preslia. 69, 129–159.

Mandák B, Pyek P. 2001. Fruit dispersal and seed banks in Atriplex sagittata: the role of heterocarpy. Journal of Ecology 89, 159–165.

Nguyen TP, Cueff G, Hegedus DD, et al. 2015. A role for seed storage proteins in Arabidopsis seed longevity. Journal of Experimental Botany 66, 6399–6413.

Orlovsky N, Japakova U, Zhang H, et al. 2016. Effect of salinity on seed germination, growth and ion content in dimorphic seeds of Salicornia europaea L. (Chenopodiaceae). Plant diversity 38, 183–189.

Racca S, Gras DE, Canal MV, et al. 2022. Cytochrome c and the transcription factor ABI4 establish a molecular link between mitochondria and ABA[dependent seed germination. New Phytologist 235, 1780–1795.

Redondo-Gómez S, Mateos-Naranjo E, Cambrollé J, et al. 2008. Carry-over of differential salt tolerance in plants grown from dimorphic seeds of Suaeda splendens. Annals of Botany 102, 103–112.

Sajeev N, Koornneef M, Bentsink L. 2024. A commitment for life: Decades of unraveling the molecular mechanisms behind seed dormancy and germination. The Plant Cell 36, 1358–1376.

Sendek A, Herz K, Auge H, et al. 2015. Performance and responses to competition in two congeneric annual species: does seed heteromorphism matter. Plant Biology 6, 1203–1209.

Tsaniklidis G, Dermitzaki E, Nikolopoulou AE, et al. 2015. Cotton seed storage effects on vigour and activities of NAD+-dependent isocitrate dehydrogenase, malate dehydrogenase and β-amylase in seedlings. Seed Science and Technology 43, 111–120.

Ungar IA, Khan MA. 2001. Effect of bracteoles on seed germination and dispersal of two species of Atriplex. Annals of Botany 87, 233–239.

Venable DL. 1985. Ecology of achene dimorphism in Heterotheca latifolia. II. Demoraphic variation within population. Journal of Ecology 73, 743–755.

Venable DL. 2007. Bet hedging in a guild of desert annuals. Ecology 88, 1086–1090.

Venable DL, Dyreson E, Morales E. 1995. Population dynamic consequences and evolution of seed traits of Heterosperma pinnatum (Aateraceae). American Journal of Botany 82, 410–420.

Wang L, Dong M, Huang ZY. 2010. Review of research on seed heteromorphism and its ecological significance. Chinese Journal of Plant Ecology, 34, 578–590.

Wang L, He XL, Zhao YJ, et al. 2011. Wheat vacuolar H+-ATPase subunit B cloning and its involvement in salt tolerance. Planta 234, 1–7.

Wang Y, Jiang W, Cheng J, et al. 2023. Physiological and proteomic analysis of seed germination under salt stress in mulberry. Frontiers in Bioscience-Landmark 28, 49.

Wang ZY, Chen WZ, Yang MH, et al. 2025. Haloxylon ammodendron adapts to desert environments through seed polymorphism during diaspore germination and seedling establishment. Frontiers in Plant Science 16, 1527718.

Wen PF, Chen JY, Wan SB, et al. 2008. Salicylic acid activates phenylalanine ammonia-lyase in grape berry in response to high temperature stress. Plant Growth Regulation 55, 1–10.

Wilkins MR, Pasquali C, Appel RD, et al. 1996. From proteins to proteomes: large scale protein identification by two-dimensional electrophoresis and arnino acid analysis. Biotechnology 14, 61–65.

Xiao J, Eziz A, Zhang H, et al. 2019. Responses of four dominant dryland plant species to climate change in the Junggar Basin, northwest China. Ecology and Evolution 9, 13596–13607.

Yan QC. 2001. Seeds. Beijing: China Agriculture Press, 261–262.

Yang TQ, Xu RH, Chen JH, et al. 2016. β-Ketoacyl-acyl carrier protein synthase I (KASI) plays crucial roles in the plant growth and fatty acids synthesis in tobacco. International Journal of Molecular Sciences 17, 1287.

Yu L, Liu Y, Peng Y, et al. 2021. Overexpression of cyanoalanine synthase 1 improves germinability of tobacco seeds under salt stress conditions. Environmental and Experimental Botany 182, 104332.

Zeng YJ, Wang YR, Zhuang GH, et al. 2004. Seed germination responses of Reaumuria soongorica and Zygophyllum xanthoxylum to drought stress and sowing depth. Acta Ecological Sinica 24, 1629–1634.

Zhang LH, Liu C, Qiu HL, et al. 2013. Cloning and function analysis of SiLEA14 from Saussurea involucrata Kar. et Kir. Acta Botanica Boreali-Occidentalia Sinica 33, 1071–1078.

Zhang MJ, Sha W, Liu B, et al. 2014. Cloning and expression analysis of glyceraldehyde-3-phosphate dehydrogenase gene RjGAPDH in Racomitrium japonicum. Journal of Anhui Agricultural Science 42, 8506–8510.

Zhumabekova Z, Xu XW, Wang YD, et al. 2020. Effects of sodium chloride and sodium sulfate on Haloxylon ammodendron seed germination. Sustainability 12, 4927.

